# Sex-specific effects of early-life adversity on adult fitness in a wild mammal

**DOI:** 10.1101/2024.08.05.606598

**Authors:** Elizabeth D. Drake, Sanjana Ravindran, Xavier Bal, Jill G. Pilkington, Josephine M. Pemberton, Daniel H. Nussey, Hannah Froy

**Author notes:** **Corresponding author:** Elizabeth Drake; Address: Institute of Ecology and Evolution, University of Edinburgh, School of Biological Sciences, Edinburgh EH9 3FL, UK.

## Abstract

Early-life adversity influences adult fitness across a range of vertebrates. In polygynous systems with intense intrasexual competition, males may be more sensitive to conditions experienced during growth and development. However, the relative importance of different aspects of the early environment and how their effects differ between the sexes remain poorly understood. Here, we used a long-term study of wild Soay sheep to characterise the early-life environment in terms of weather, infection, resource competition and maternal investment, and test the hypothesis that males are more vulnerable to early adversity than females. Birth weight positively predicted lifetime breeding success in both sexes, suggesting a classic ‘silver spoon’ effect, though the effects were stronger in males. Males experiencing high population densities in their first year had lower lifetime breeding success suggesting lasting negative consequences of nutritional stress, but there was no association in females. In contrast, challenging weather in the first winter of life appeared to act as a selective ‘filter’, with males surviving these harsh conditions having higher adult fitness. Our findings further evidence the important long-term fitness consequences of early-life adversity in wild vertebrates, highlighting that different aspects of the early environment may shape later fitness in different and sex-specific ways.

## Introduction

Development is particularly sensitive to the environment, and the conditions experienced during this life stage can have lasting impacts on physiological function, demographic rates and ultimately lifetime fitness [1,2]. Thus, early-life environment can have important consequences for ecological and evolutionary processes, shaping population dynamics and life history evolution [2–4]. Numerous studies show that favourable conditions during early life can have positive consequences for adult body size, reproduction and survival, coined ‘silver spoon’ effects [5–8]. Similarly, many human and wild primate studies have developed composite, cumulative measures of early-life adversity and shown that high adversity scores predict lower adult health and fitness [9–11]. As well as directly impacting development, challenging early conditions may act as a selective ‘filter’, removing weak individuals from a cohort and leaving a group of adults that is fitter on average than cohorts experiencing benign conditions [12–15]. Natural populations experience complex, multivariate environments involving diverse challenges including resource competition, harsh weather, constraints on parental investment and parasite infection [1]. These challenges could impact development and fitness via different processes and to different degrees, yet most studies to date have used composite measures of adversity or a small number of environmental variables [12,16–18]. In this study, we provide a rare test of how different aspects of early-life adversity shape reproductive performance, survival and lifetime fitness in a wild vertebrate population living in a highly variable environment.

In polygynous systems, the effects of early-life conditions on later-life fitness are expected to differ between the sexes due to the different selection pressures on males and females [2,19,20]. Males may have greater nutritional and energy requirements and increased sensitivity to environmental adversity during gestation and neonatal periods due to their more rapid growth rates, larger body size and the costs of developing secondary sexual characteristics [21]. This heightened sensitivity may mean males in sexually dimorphic species are more adversely affected by poor conditions in early life [14,22]. This may lead to sex-biases in juvenile mortality which could impose a stronger selective filter on males, as well as increasing the likelihood of showing long-lasting silver spoon effects [4]. However, studies that have explicitly tested for differing consequences of early-life adversity between the sexes have found mixed results, and few have done so in polygynous mammals [15,23–25]. To thoroughly examine the connections between the early and later stages of life, it is essential to utilize individual-based data spanning entire lifespans, from a system where high-quality data are available for both sexes. Here, we use such data from a long-term study of a highly polygynous mammal to test the prediction that early-life adversity has a stronger effect on adult fitness in males than females.

The long-term study of Soay sheep on St Kilda offers detailed, longitudinal data on morphology, life history and demography collected across the lifetimes of thousands of individuals over four decades [26]. The early-life environmental factors impacting first year survival are well understood in this system [21,27–31]. Experience during the first year of life of higher population density, litter size and parasite burden, lower body weight and wetter and windier winter weather conditions during the first year of life are all known to decrease lamb winter survival [26,29,30,32]. Increased natal litter size and higher parasite loads are also associated with reduced reproductive success [26,33,34]. Additionally, the mating system is highly polygynous, and males attain considerably larger adult size than females [26]. Adult annual winter mortality is higher and more sensitive to resource competition in the preceding year in males compared to females [28]. In this study, we link this broad suite of early environmental challenges to adult fitness in both sexes, and test whether effects are stronger in males than females. We demonstrate that different aspects of the environment shape adult fitness through different processes (i.e., silver spoon versus filtering effects) and through different components of fitness (i.e., reproduction versus survival), highlighting the complexity of sex-dependent early-life effects.

## Methods

### Study system and data

Since 1985, the Soay sheep resident to the Village Bay area of Hirta, St. Kilda, Scotland have been the subject of an individual-based study [26]. During the mating season in October-November, males compete to gain access to oestrous ewes and sequentially mate with multiple females while forming consort pairs. After an approximately five-month gestation period, lambs are born April-May and >90% of lambs born in the study area are caught within a few weeks of birth. At capture, lambs are uniquely marked with ear tags, tissue sampled and weighed. Each August, 50-70% of the population are caught, weighed and faecal samples are taken. Regular censuses mean that the fate of individuals is known with a high degree of certainty. Mortality peaks in late winter/early spring, and around 80% of deceased sheep are found [26].

In this study, we used data collected between 1985 and 2022. To test for persistent effects of early-life adversity on fitness, we developed a list of candidate early-life metrics and tested whether they independently predicted first year survival. We then tested whether they were associated with lifetime fitness (see below) in individuals that survived beyond their first year of life, and whether those associations differed between the sexes.

### Early-life measures

Natal litter size: Coded as singleton or twin. Around 18% of Soay sheep births are of twins, and triplets are very rare (7 sets recorded, grouped with twins in this analysis). Twins are lighter than singletons at birth and in August and have reduced first year survival [30].

Maternal loss: A binary trait indicating whether an individual lost their mother in their first year of life (i.e. whether mother died before 1 May in the year following birth or survived). For a very small number of cases, the mother’s month of death was not known with certainty. In these cases, we assumed the mother had died over winter in the recorded year of death (<1% of records) or, if year of death was unknown, we assumed the mother had died over winter in the year she was last seen in a census (<1%).

Winter NAO in gestational winter and in first winter: NAO is a measure of atmospheric pressure across the North Atlantic region, based on the relative changes in pressure between stations in Portugal and Iceland. Positive NAO values indicate mild, stormy and wet winter conditions in Northern Europe and negative values indicate cold, calm and dry winter weather. Positive winter NAO values predict increased winter mortality in this population [28]. We tested effects of the NAO both the winter before (gestational winter) and the winter after birth (first winter) using December-March Hurrell North Atlantic Oscillation Index (station-based) from the NCAR Climate Data Guide (https://climatedataguide.ucar.edu/climate-data/hurrell-north-atlantic-oscillation-nao-index-station-based).

Population size in year of birth: Birth and death dates were used alongside census records to estimate the total number of individuals alive resident in the Village Bay area on 1^st^ October each year. High population sizes are associated with increased subsequent winter mortality, particularly in lambs, presumably through increased resource competition [28,31].

Parasite exposure in year of birth: Since 1988, the number of Strongyle nematode eggs has been counted as a proxy of parasite burden in each faecal sample collected from sheep at August capture (faecal egg counts, FEC). Counts were made using a modified McMaster technique [35]. Strongyle gastrointestinal parasites are highly prevalent in this population and FEC predicts summer weight and overwinter mortality in lambs [31,34,36]. Here, we used the average FEC across a cohort of lambs in August as an indicator of parasite exposure (following [37]). Two extreme outlying individual values (> 10,000) were removed before calculating the mean for each cohort. FEC data were unavailable for the earliest cohorts (1985-1987) and for >90% lambs in 2002, and counts were performed using a different method in recent years (2019-2021). See the ‘Statistical analyses’ section below for how these missing data were handled.

Birth weight: Calculated in kilograms for individuals caught within the first seven days of life (at least 60% of individuals in each cohort). Birth weight has previously been positively associated with subsequent growth rates and first winter survival in this system [32]. Variation in age at capture was accounted for by taking the residuals of a generalised additive model of birth weight with a smoothing term for age at capture in days (package *mcgv*; [38]). An outbreak of foot and mouth in 2001 and the Covid-19 pandemic in 2020 meant none and 23% of these cohorts were captured and weighed within a week of birth, respectively. See below for how missing data (from the 2001 and 2020 cohorts, and where individuals were not caught within the first week of life) were handled.

The correlations among early-life measures are shown in Figure S1 and Table S1. Collinearity between fixed effects was tested for each model using the package performance [39]; all variance inflation factors were less than 3, suggesting little collinearity between covariates.

### Fitness components

First year survival was defined as survival to 1 May in the year following birth (0: died before 1 May in year following birth; 1: survived to 1 May in year following birth). A small proportion of individuals were assigned assuming late winter/early spring mortality in the recorded year of death when death month was unknown (<5%) or based on the last time they were seen in a census (<1%). Our analyses of first year survival included data from all individuals born between 1985 and 2021 who survived at least past 1 October in their year of birth where sex, birth year, maternal identity and the fate of the mother were known (N=5,073).

Our analyses of adult fitness components were restricted to individuals who survived beyond their first year of life, to ensure any associations with early-life adversity were independent of first year survival. Reproductive performance in both sexes was calculated based on a multigenerational genetic pedigree, inferred using a subset of 431 unlinked Single Nucleotide Polymorphisms (SNPs) derived from the Illumina Ovine 50K SNP array using the R package Sequoia [39,40]. For a small number of cases where SNP genotype information was not available, parentage assignments were made using field observations (for females) or from microsatellite data [41].

Lifetime breeding success (LBS) was defined as the number of offspring born to or sired by an individual over their lifespan. This ranged between 0-20 for females and 0-94 for males. We also considered different components of LBS: longevity and annual reproductive performance. Longevity was the age of an individual at death, ranging from 1-15 for females and 1-11 for males. Our analyses of LBS and longevity were restricted to individuals born 1985-2017 that had completed their natural lifespans and for whom year of death was known (N=1721; 1034 females and 687 males). Where information on the month of death was missing (∼12% of individuals), late winter/early spring mortality was assumed when assigning longevity unless an individual was seen in a census later that year.

Analyses of annual reproductive performance measures included individuals that were still alive. We used all observations from individuals born 1985-2021 where age ≥ 1. Breeding probability was a binary trait indicating whether an individual did (1) or didn’t (0) give birth to or sire an offspring in each year of their life (N=9,572 observations of 2,232 individuals). For females that bred in a given year, twinning probability was a binary trait indicating whether the female gave birth to a single lamb (0) or twins (1) (N=5,580 observations of 1,092 females). For males that bred in a given year, offspring number was the number of live offspring they sired that year, ranging from 1–22 (N=884 observations of 378 males).

### Statistical analyses

We tested for associations between our metrics of early-life adversity and our fitness components using generalized linear mixed effects models (GLMMs). First year survival, breeding probability and female twinning probability were modelled with a binomial error distribution and logit link function. Lifetime breeding success, longevity and male offspring number were modelled with a negative binomial error distribution and log link function.

First winter NAO, population size in year of birth, mean cohort FEC and birth weight were included in all models as fixed covariates. Natal litter size and maternal loss were included as two-level fixed factors. Gestational winter NAO was additionally included as a covariate in the first-year survival model, but since it did not independently predict first year survival it was omitted from the analyses of adult fitness (see Results). Sex was included as a two-level fixed factor in all models that included both males and females to account for differences in average trait values between the sexes. To test for sex-specific effects of early-life adversity, we also included two-way interactions between sex and each of our early-life metrics in these models.

All models included cohort and maternal identity as random intercept terms to account for shared environmental conditions experienced over the lifetime and non-independence of offspring born to the same mother, respectively. The models of annual fitness components (breeding probability, female twinning probability and male offspring number) also included measurement year and individual identity as random intercept terms. The annual fitness models additionally included a quadratic effect of age in years as a covariate to account for age-related variation in reproductive performance [42]. Interactions between the age terms and sex were included in the model of breeding probability to allow the ageing trajectories to differ between males and females.

All covariates were z-standardized prior to inclusion in the models to aid interpretability (mean=0 and standard deviation=1). Interactions that were not statistically significant at the P>0.05 level were sequentially deleted from the model, beginning with the interaction with the highest p-value. All main effects were retained in the model regardless of statistical significance (see Table S2 for full models including all interaction terms).

Where data on cohort mean faecal egg count or birth weight were missing (∼15% and ∼20% of lambs, respectively), we used a data imputation approach assuming the data were missing at random to avoid a reduction in sample size [43]. For each of our models, the relevant data subset was used to calculate the mean value for each trait (cohort mean faecal egg count or birth weight), and this was used where data were missing. To check whether this approach impacted our results, we re-ran each of our models excluding the imputed data, and examined whether the results remained the same for cohort mean faecal egg count or birth weight, respectively. Our results remained unchanged (data not shown).

Lastly, we re-ran our final models controlling for mean adult weight to test whether the associations between early-life environment and adult fitness were mediated by adult body weight. Adult weight in kilograms was measured in August. To control for variation in capture age and sex, we took the residuals of a generalised additive model of August weight with a smoothing term for age at capture (in years), a two-level fixed factor for sex, and the interaction between the two (*mgcv*; [38]). We then calculated the mean residual August weight for each individual from all captures where age ≥ 1 and included this as a fixed covariate in our models of adult fitness.

All statistical analysis in this work were performed using R and RStudio [44,45]. All models were run using the package *glmmTMB* [46], in which significance of fixed effects was determined using Wald chi-square tests. Variance and 95% confidence intervals for random effects were calculated using the package *stats* [45]. *ggplot2* [47] and *ggeffects* [48] packages were used for plotting.

## Results

All our measures of early-life adversity were independently associated with first year survival, with the exception of gestational winter NAO (Table 1, Figure 1). Twins and individuals who lost their mothers during their first year were less likely to survive their first year of life, as were lambs who were born lighter (Table 1, Figure 1A, 1B & 1G). Those born in years with higher population sizes were also less likely to survive their first year, and this association was stronger for male than female lambs (significant sex-by-population size interaction in Table 1, Figure 1C). Lambs who experienced harsh weather conditions during their first winter (high NAO values) had lower survival probabilities, and this association was again stronger for male than female lambs (Table 1, Figure 1D). Similarly, lambs in cohorts with higher mean FEC were less likely to survive to the following spring, and the association was stronger for males (Table 1, Figure 1F).

**Table 1.** Fixed and random effect estimates from a binomial GLMM of first year survival in Soay sheep. Statistically significant fixed effects are highlighted in bold. The reference levels for the fixed factors were: sex (female), natal litter size (singleton) and maternal loss (mother died in first year of life). The fixed covariates were scaled to mean=0 and standard deviation=1.

| N=5073 |  |  |  |  |  |  |
| --- | --- | --- | --- | --- | --- | --- |
| Fixed effects | Estimate | Std.<br>Error | p-value | Random effects | Variance | 95% CI |
| **Intercept** | **-0.631** | **0.189** | **0.001** | Maternal identity | 0.548 | 0.381-0.788 |
| **Sex (male)** | **-0.845** | **0.083** | **<0.001** | Birth year | 0.717 | 0.430-1.234 |
| **Natal litter size (twin)** | **-0.723** | **0.118** | **<0.001** |  |  |  |
| **Maternal loss (mother survived)** | **0.803** | **0.124** | **<0.001** |  |  |  |
| **Population size YOB** | **-0.828** | **0.201** | **<0.001** |  |  |  |
| Gestational winter NAO | 0.177 | 0.179 | 0.323 |  |  |  |
| **First winter NAO** | **-0.561** | **0.153** | **<0.001** |  |  |  |
| **Cohort mean FEC** | **-0.459** | **0.182** | **0.012** |  |  |  |
| **Birth weight** | **0.322** | **0.046** | **<0.001** |  |  |  |
| **Sex (male): Population size YOB** | **-0.292** | **0.108** | **0.007** |  |  |  |
| **Sex (male):First winter NAO** | **-0.302** | **0.084** | **<0.001** |  |  |  |
| **Sex (male):Cohort mean FEC** | **-0.236** | **0.100** | **0.018** |  |  |  |

**Figure 1.**
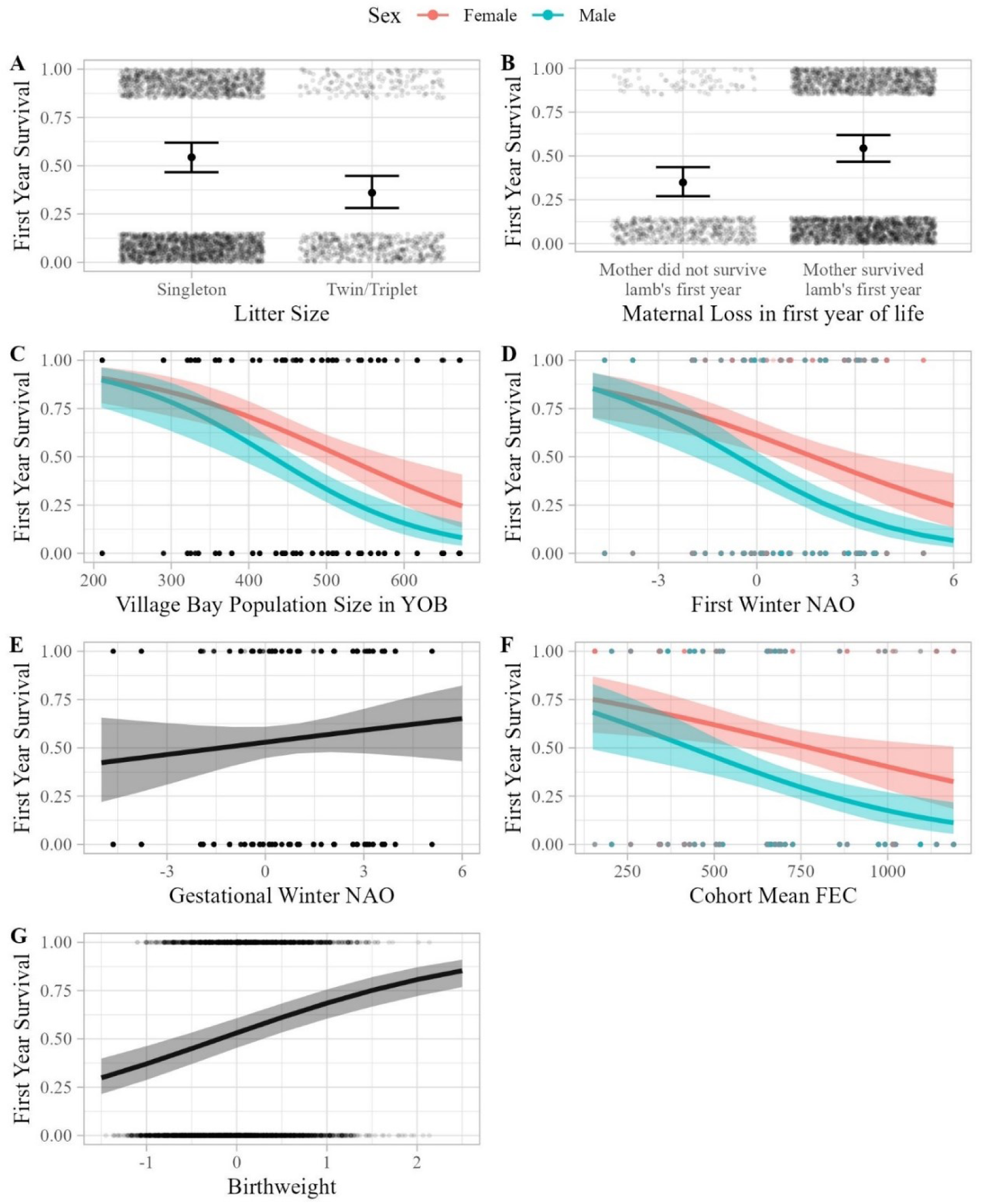
Estimates of the effects of different measures of early-life adversity on first year survival in Soay sheep (N=5074 observations). Plots show independent effects of (A) natal litter size, (B) maternal loss during the first year of life, (C) gestational winter NAO, (D) first winter NAO, (E) cohort mean FEC in year of birth, (F) population size in year of birth and (G) birth weight. Points show the (jittered) raw data. Lines and shading show the predictions and associated 95% CIs from the model in Table 1, where all other covariates were held at the mean and factors at the reference level, except maternal loss, which was held at ‘mother survived first year of life’. Plots with coloured lines indicate where the association differed between the sexes; females shown in red, males in blue. All estimates were statistically significant except for gestational winter NAO.

The lifetime breeding success (LBS) of individuals that survived beyond their first year was associated with measures of early-life adversity, sometimes in sex-specific ways (Table 2, Figure 2). Maternal loss in the first year of life was associated with reduced LBS in both sexes (Table 2, Figure 2B). Individuals that were born lighter also had lower LBS, but the effect was stronger in males (Table 2, Figure 2F). There was a significant interaction between sex and population size in the year of birth: males born in high density years had lower LBS, whereas there was no association in females (Table 2, Figure 2C). First winter NAO was also positively associated with LBS in males but not females: males who experienced harsher weather conditions during their first winter performed better as adults (Table 2, Figure 2D). Natal litter size and cohort mean FEC were not significantly associated with LBS in adults (Table 2; Figure 2A & 2E).

**Table 2.** Fixed and random effect estimates from a negative binomial GLMM of lifetime breeding success in Soay sheep that survived beyond the first year of life. Statistically significant fixed effects are highlighted in bold. The reference levels for the fixed factors were: sex (female), natal litter size (singleton) and maternal loss (mother died in first year of life). The fixed covariates were scaled to mean=0 and standard deviation=1. Dispersion parameter for negative binomial model 0.932.

| N=1721 |  |  |  |  |  |  |
| --- | --- | --- | --- | --- | --- | --- |
| Fixed effects | Estimate | Std.<br>Error | p-value | Random effects | Variance | 95%CI |
| **Intercept** | **1.305** | **0.150** | **<0.001** | Maternal identity | 0.209 | 0.141-0.310 |
| **Sex (male)** | **-0.723** | **0.073** | **<0.001** | Birth year | 0.128 | 0.061-0.265 |
| Natal litter size (twin) | 0.059 | 0.103 | 0.570 |  |  |  |
| **Maternal loss (mother survived)** | **0.271** | **0.136** | **0.046** |  |  |  |
| Population size YOB | 0.049 | 0.092 | 0.593 |  |  |  |
| First winter NAO | 0.017 | 0.079 | 0.828 |  |  |  |
| Cohort mean FEC | -0.027 | 0.079 | 0.732 |  |  |  |
| **Birth weight** | **0.159** | **0.044** | **<0.001** |  |  |  |
| **Sex (male):Population size YOB** | **-0.222** | **0.069** | **0.001** |  |  |  |
| **Sex (male):First winter NAO** | **0.157** | **0.068** | **0.021** |  |  |  |
| **Sex (male):Birth weight** | **0.161** | **0.068** | **0.017** |  |  |  |

**Figure 2.**
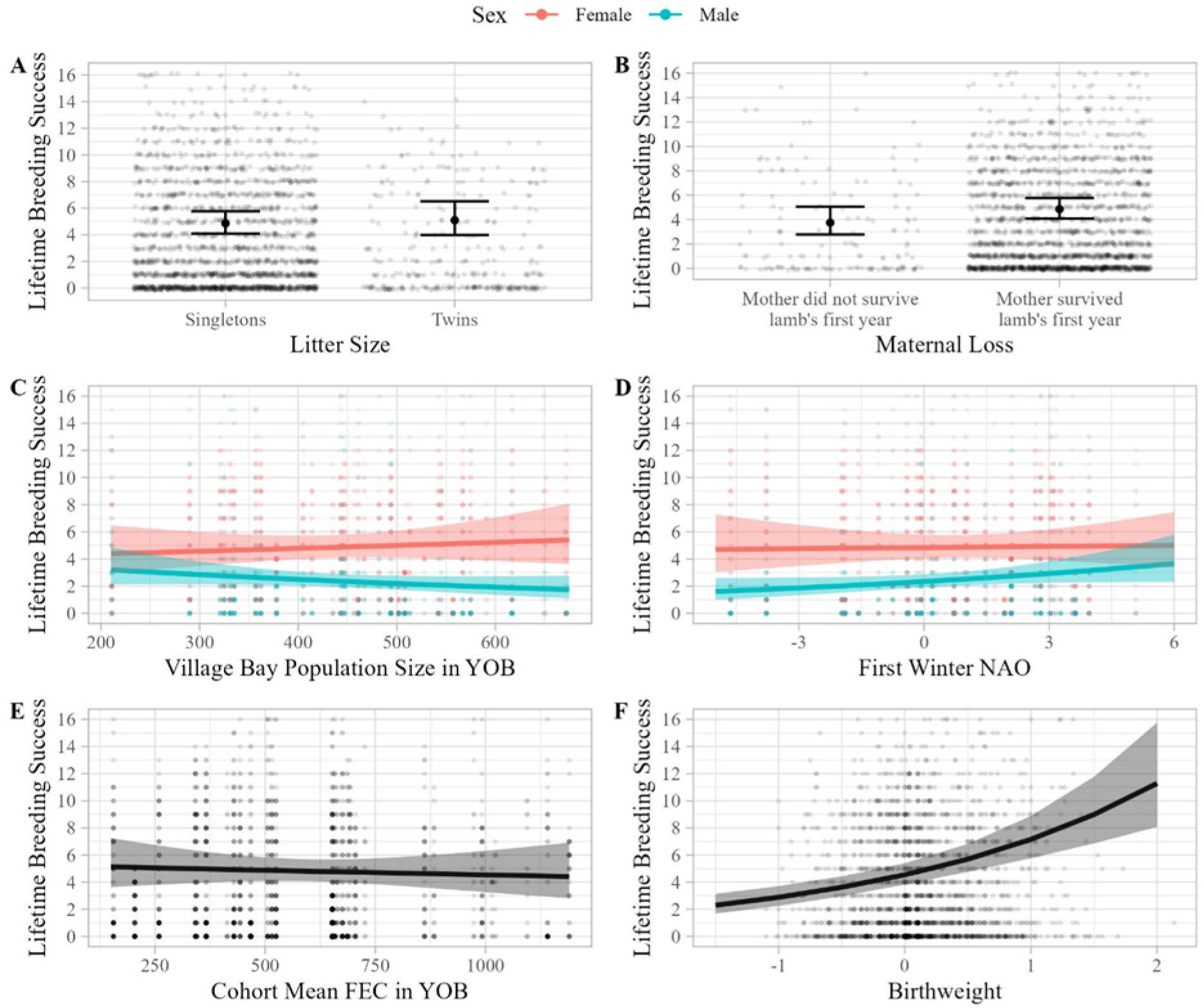
Estimates of the independent effects of (A) natal litter size, (B) maternal loss during the first year of life, (C) first winter NAO, (D) cohort mean FEC in year of birth (E) population size in year of birth and (F) birth weight on lifetime breeding success in Soay sheep that survived beyond the first year of life (N=1721 observations). Points show the (jittered) raw data. y-axes are limited to LBS < 17 for clarity (representing > 97% of observations). Lines and shading show the predictions and associated 95% CIs from the model in Table 2, where all other covariates were held at the mean and factors at the reference level, except maternal loss, which was held at ‘mother survived first year of life’. Plots with coloured lines indicate where the association differed between the sexes; females shown in red, males in blue. The effect of birth weight was statistically significant, as were the effects of population size in year of birth and first winter NAO in males.

Adult longevity was positively associated with birth weight in both sexes (estimate=0.063 ±0.020SE, p=0.001; Table S3, Figure 3B). There was a significant interaction between the effects of sex and first winter NAO; males who experienced harsh winter weather during their first year had longer lifespans, whereas female longevity showed no association with first winter NAO (estimate in females=0.041 ±0.053, p=0.441; estimate in males vs females=0.120 ±0.036, p=0.001; Table S3, Figure 3A). There was no significant association between litter size, maternal loss, population size in year of birth or cohort mean FEC and adult longevity (Table S3).

**Figure 3.**
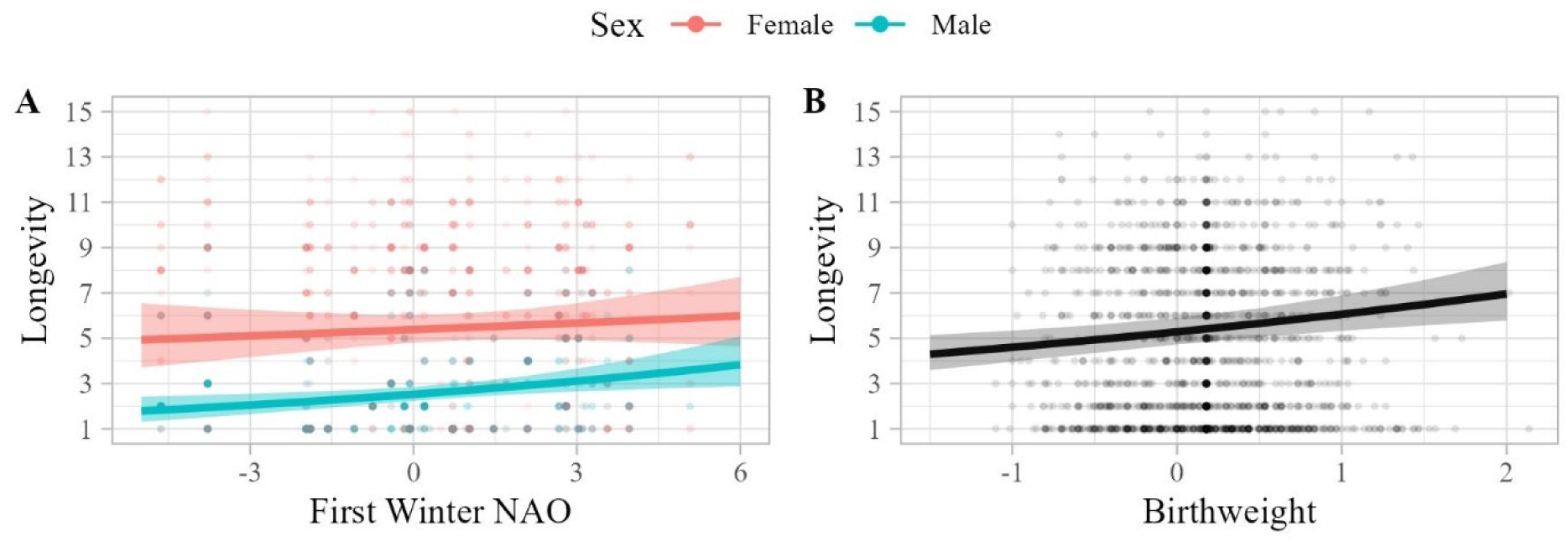
Estimates of the statistically significant, independent effects of (A) first winter NAO and (B) birth weight on longevity in Soay sheep that survived beyond the first year of life (N=1721 observations). Points show the raw data. Lines and shading show the predictions and associated 95% CIs from the model in Table S3 where all other covariates were held at the mean and factors at the reference level, except maternal loss, which was held at ‘mother survived first year of life’. The plot with coloured lines indicates where the association differed between the sexes; females shown in red, males in blue.

Annual breeding probability was positively associated with birth weight in both females and males (estimate=0.260 ±0.054, p<0.001; Table S4, Figure 4C). Maternal loss in the first year of life was also significantly associated with reduced adult breeding probability in both sexes (effect of mother surviving=0.488 ±0.190, p=0.010; Table S4, Figure 4A). Although higher population sizes in the year of birth were associated with decreased annual breeding probability for both sexes, the effect of population size was significantly stronger in males than females (estimate in females=-0.301 ±0.111, p=0.007; estimate in males vs females=-0.350 ±0.121, p=0.004; Table S4, Figure 4B). The association between annual breeding success and cohort mean FEC differed between the sexes, but the association was not statistically significantly different from zero in either males or females (estimate in females=-0.056 ±0.101, p=0.575; estimate in males=0.227 ±0.136, p=0.089; Table S4). Litter size and first winter NAO were not significantly associated with annual breeding probability (Table S4).

**Figure 4.**
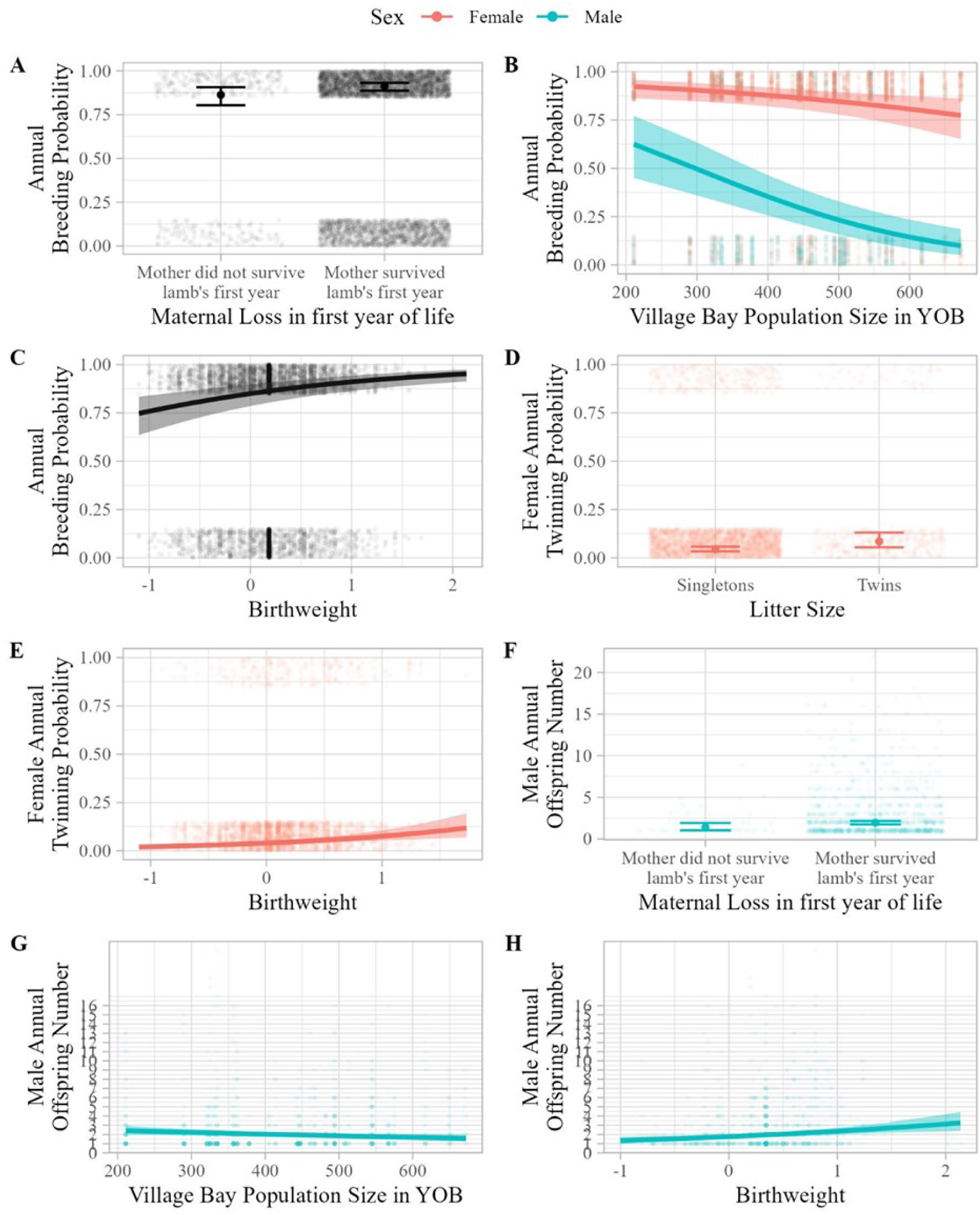
Estimates of the statistically significant effects of (A) maternal loss in the first year of life, (B) population size in year of birth and (C) birth weight on annual breeding probability; (D) natal litter size and (E) birth weight on female annual twinning probability; and F) population size in year of birth and (G) birth weight on male annual offspring number. Points show the raw (jittered) data. Lines and shading show the predictions and associated 95% CIs from the models in Table S4, S5 & S6 where all other model covariates were held at the covariate mean and factors at the reference level, except maternal loss, which was held at ‘mother survived first year of life’. Effects in females shown in red and males in blue.

Among females that successfully reproduced, females who were heavier at birth were more likely to produce twins (estimate=0.301 ±0.081, p<0.001; Table S5, Figure 4E). Annual twinning probability was also significantly higher in females who were twins themselves (estimate=0.702 ±0.246, p=0.004; Table S5, Figure 4D). There was no significant association between maternal loss, population size in year of birth, first winter NAO or mean cohort FEC and annual twinning probability (Table S5). There was a positive association between birth weight and offspring number in males that successfully reproduced, with males who were born heavier siring significantly more offspring each year (estimate=0.125 ±0.038, p=0.001; Table S6, Figure 4H). Males who lost their mother in their first year of life had fewer offspring annually (effect of mother surviving=0.334 ±0.163, p=0.040; Table S6, Figure 4F. Population size in the year of birth was negatively associated with male offspring number, with males born in high density years siring fewer offspring annually (estimate=-0.104 ±0.048, p=0.030; Table S6, Figure 4G). Litter size, first winter NAO and cohort mean FEC were not significantly associated with annual male offspring number (Table S6).

The positive associations between birth weight and male lifetime breeding success, breeding probability and male offspring number remained statistically significant when controlling for the effects of mean adult weight, though the effect sizes were somewhat reduced (Table S7A, S7C & S7E). In contrast, birth weight was no longer positively associated with female lifetime breeding success, longevity or female twinning probability once mean adult weight was accounted for (Table S7A, S7B & 7D). Once mean adult weight was included, the interaction between sex and population size in the year of birth was no longer significant in the models of LBS and breeding probability, and there was no negative association between population size and male offspring number (Table S7A, S7C & S7E). The associations between first winter NAO and male LBS and male longevity remained when accounting for mean adult weight (Tables S7A & S7B).

## Discussion

Our results add to the growing literature demonstrating that early-life environmental conditions have important, long-term effects on adult fitness in wild vertebrate populations [1,17,49–51]. The detailed, longitudinal data collected over four decades on St Kilda enabled us to dissect the effects of multiple aspects of the early-life environment on adult fitness in both male and female Soay sheep. We found that different components of the environment experienced during the first year of life impacted adult fitness independently, and in strikingly different ways. Birth weight was positively associated with adult lifetime breeding success (LBS) in both sexes, but the effect was stronger in males and acted via effects on both longevity and breeding success. However, the negative effect of population size in the year of birth on adult LBS was only detected in males, and operated through effects on reproductive performance but not longevity [52]. These findings are consistent with so-called ‘silver spoon’ effects: individuals that experienced favourable conditions during development performed better as adults [1,6,51]. In contrast, weather conditions experienced during the first winter were predictive of adult male LBS, but the effect was in the opposite direction. Although wet, windy first winters were associated with increased lamb mortality, those males that survived these harsh winters had higher adult LBS than those that experienced more favourable cold, dry winters. This was driven by an effect on adult longevity, and is consistent with a selective ‘filter’ effect in which only more robust, higher quality individuals survive to adulthood [12–14]. These marked differences in the relationships between early-life environment, sex and adult fitness highlight the importance of understanding the detail of the processes linking different aspects of early-life adversity and later health and fitness.

First-year survival was independently predicted by multiple components of the early-life environment: twins, lighter born lambs, those that lost their mothers in the first year of life, as well as those experiencing higher cohort mean parasite burdens, higher population densities, and harsher winter weather were less likely to survive their first winter, confirming previous findings in this study population [27–31]. The associations with cohort mean FEC, population size in the first year of life and first winter NAO were stronger in male than female lambs. This supports our prediction that males would be more sensitive to early-life conditions owing to their larger size and more rapid growth trajectories [21]. Effects of maternal mortality over the first year on lamb survival have not previously been investigated in Soay sheep, although other vertebrate studies have shown that ‘orphaning’ impacts offspring survival and later fitness [20,53–55]. Although most lambs are weaned by late summer, long before most maternal mortality occurs, they remain closely associated with their mothers for some time, and this effect could reflect social or physiological costs of maternal mortality for the lamb. However, this could also be driven by other environmental factors jointly shaping mother and offspring survival not captured by other early-life variables in our models. Previous studies have documented a relationship between lamb survival and maternal age, with lamb survival lowest in the final year of a female’s life, which is consistent with the relationship observed here [29]. The weak but significant relationship between maternal loss in early life and adult lifetime breeding success (Figure 2B; Table S3) does suggest there may be long-term effects of orphaning in this system and merits further, in-depth study.

Birth weight was positively associated with multiple components of fitness in both male and female sheep that survived beyond their first winter (Figures 2F, 3B, 4C, 4E&4G). Birth weight has been shown to affect later-life reproduction and survival in natural, domestic and laboratory populations [56–58], consistent with ‘silver spoon’ effects of early environment. Unlike our annual environmental variables, birth weight varies within years and reflects variation among individual mothers in their ability to care for, and invest in, their offspring [30]. In Soay sheep, birth weight is influenced by maternal genetic and environmental effects, and is positively associated with lamb weight in summer and adult weight [26,32,39,59,60]. Birth weight was predictive of male LBS independent of mean adult weight, suggesting that the effects are not solely due to heavier neonate lambs growing faster and developing into heavier, higher condition adults. The observed effects of birth weight may reflect long-term impacts on development and physiological function of maternal investment during both gestation and lactation, as mothers in better condition are likely to invest more during both phases of development. They could also reflect fine-scale spatial variation in the mother’s environment as home range quality has been shown to influence fitness, and this is shared with the offspring over the first year of life [61]. The effect of birth weight on LBS was stronger in males than females, in line with our predictions and previous studies of wild mammals, which suggest that males gain greater fitness benefits from being born heavier than females [56]. Although female LBS was associated with birth weight, this association was driven by the fact that heavier born females were heavier as adults. In contrast, the associations between birth weight and multiple components of fitness were independent of adult weight in males, suggesting that early nutritional deficits may have more fundamental consequences. Our findings highlight the sensitivity of the earliest stages of development and demonstrate that compensatory growth is not always sufficient to overcome nutritional stress experienced very early in life.

First winter NAO had an unexpected positive effect on the longevity of males that survived challenging conditions during their first winter, which translated into an increase in male adult LBS (Figures 2D & 3A). Winter NAO is thought to predict fitness in the sheep because it captures the thermoregulatory challenges associated with the frequent and severe winter storms that kill already food-limited sheep on St Kilda [28,31,62]. It is therefore an indicator of a direct environmental driver of lamb mortality in late winter that may act as a selective ‘filter’, removing weak individuals and leaving only more robust individuals in these cohorts. Long-term studies in other systems have shown that strong early-life viability selection can produce adult cohorts with increased longevity [12–14,19]. Such cohort filtering effects may be expected to manifest through longevity rather than reproduction as they operate through differences in average physiological condition and resilience among birth cohorts, which could drive differences in adult survival. The effects of first winter NAO were sex-specific; there were no significant effects of first winter weather on female adult fitness traits (Figure 2D&3A). This fits with both the general prediction that viability selection is stronger in males in polygynous systems [63,64], and the finding that male lamb survival is more strongly influenced by winter NAO in this population (Figure 1D).

We also found long-term, sex-specific effects of population size in the year of birth on adult LBS (Figure 2C). Population size did not influence adult longevity, instead depressing annual breeding probability in both sexes, and the number of offspring sired in males (Figures 4B&4F). These ‘silver spoon’ effects contrast with the findings for first winter NAO. Their opposing effects on adult LBS likely reflects the fact that these two early-life predictors influence adult fitness via different processes. Annual population size shapes the development and demography of the sheep via effects on resource availability and density dependence.

Rather than reflecting a direct mortality risk in winter, it captures the level of competition for food and mates across the study area over the entire year. Other studies have shown similar negative effects of high early-life population density on later fitness in wild vertebrates [7,65,66]. These are typically attributed to reduced resource availability and nutritional stress in early life having a negative impact on development and thus adult condition and fitness [19,49]. On St Kilda, this effect may be most acutely felt during the lamb’s first autumn and winter when the animals are growing fast, and plant food resources are at their most limited. This could be somewhat independent of birth weight effects, if these are more reflective of maternal investment and conditions experienced through the previous winter and spring. High sheep numbers, reflecting high competition for limited food late in the year, should negatively impact growth rates and, potentially adult size and condition. This would fit with the finding that early population size effects are stronger for males, for whom growth is more rapid from the first year onwards, and adult fecundity is more strongly size- and condition-dependent compared to females [26,52]. Accounting for mean adult body weight rendered the sex-by-population size interaction non-significant in the models of LBS and annual breeding probability, and the effect of population size non-significant in the model of male offspring number (Table S7). This supports the hypothesis that first year population size effects on reproductive performance are mediated by resource limitation and competition impacting growth and ultimately adult body size and condition.

Early-life environmental conditions influence the developmental trajectories, behaviour, life history and ultimately fitness of individuals, and can impose strong selective filters on cohorts, in natural populations [1,12,67]. Although there is abundant evidence to support the existence and ecological and evolutionary significance of such effects, natural populations experience a range of different environmental challenges which may not all have the same effect within a given population. Human and primate studies have sought to combine and simplify multiple early-life environmental factors into single metrics of cumulative early-life adversity [9–11], although a recent study of wild hyaenas showed that this approach reduced the power to explain variation in adult fitness [17]. Our detailed analysis of the long-term fitness effects of well-understood early environmental measures demonstrates that different aspects of the early environment can influence adult fitness via different processes (selective filtering or silver spoon effects), different fitness components (longevity or survival), and in sex-specific ways. Our findings highlight both the importance of long-term, individual-based studies for our understanding of variation in fitness and demography in wild animal systems [68] and the need for a more detailed understanding of how specific environmental pressures in early life shape development, viability selection and adult reproduction, health and survival.

## Acknowledgements

We are grateful to everyone who has been involved in the long-term study of Soay sheep on St Kilda, Scotland over the last 39 years, particularly Ian Stevenson, Michael Morrisey, Jon Slate, Susan Johnston, Tim Clutton-Brock, Loeske Kruuk and the many field volunteers. We thank the funders of the long-term project: the Natural Environment Research Council, the Biological and Biotechnology Research Council and the European Research Council. We are also grateful to the National Trust for Scotland for permissions to work on St Kilda, and QinetiQ and Kilda Cruises for logistical assistance in the field. EDD was funded by the NERC E4 Doctoral Training Partnership and HF by a Royal Society University Research Fellowship.

## Data availability statement

All data used in this manuscript will be made available on on-line upon final acceptance of the manuscript

## Statement of authorship

EDD, HF, DHN and SR designed the study; JGP, JMP and XB led the fieldwork and data collection; and EDD and HF performed analyses with input from SR. EDD produced the figures and wrote first draft of manuscript with input from HF and DHN; all authors contributed to revisions.

The authors declare no conflict of interest.

