## Supplemental Material for "Sex-specific effects of early-life adversity on adult fitness in a wild mammal"

Table S1. Pearson's moment correlation coefficients and p-values for the relationships between the early life measures. Significant correlations are highlighted in bold.

| | Maternal loss | First winter NAO | Birth weight | Cohort mean FEC | Population size YOB | Natal litter size |
|---|---|---|---|---|---|---|
| **Gestational winter NAO** | **0.116 p < 0.001** | **0.106 p < 0.001** | **-0.096 p < 0.001** | **-0.274 p < 0.001** | **-0.588 p < 0.001** | **0.026 p = 0.031** |
| **Natal litter size** | **-0.068 p < 0.001** | **0.044 p < 0.001** | **-0.327 p < 0.001** | -0.019 p = 0.113 | -0.006 p = 0.605 | |
| **Population size YOB** | **-0.198 p < 0.001** | **-0.151 p < 0.001** | **-0.069 p < 0.001** | **0.586 p < 0.001** | | |
| **Cohort mean FEC** | **-0.185 p < 0.001** | 0.017 p = 0.171 | -0.009 p = 0.522 | | | |
| **Birth weight** | 0.023 p = 0.099 | 0.006 p = 0.664 | | | | |
| **First winter NAO** | **-0.138 p < 0.001** | | | | | |

Table S2. Fixed and random effect estimates from GLMMs of A) first year survival (binomial), B) lifetime breeding success (negative binomial), C) longevity (negative binomial), and D) breeding probability (binomial) in Soay sheep. Models include two-way interactions regardless of statistical significance. Statistically significant fixed effects are highlighted in bold. The reference levels for the fixed factors were: sex (female), natal litter size (singleton) and maternal loss (mother died in first year of life). The fixed covariates were scaled to mean=0 and standard deviation=1.

| **Fixed effects** | **Estimate** | **Std. Error** | **p-value** | **Random effects** | **Variance** |
|---|---|---|---|---|---|
| A) First year survival (N=5073) | | | | | |
| **Intercept** | **-0.618** | **0.206** | **0.003** | Maternal identity | 0.553 |
| **Sex (male)** | **-0.865** | **0.245** | **<0.001** | Birth year | 0.732 |
| **Natal litter size (twin)** | **-0.860** | **0.155** | **<0.001** | | |
| **Maternal loss (mother survived)** | **0.819** | **0.155** | **<0.001** | | |
| **Population size YOB** | **-0.818** | **0.204** | **<0.001** | | |
| Gestational winter NAO | 0.198 | 0.184 | 0.281 | | |
| **First winter NAO** | **-0.562** | **0.154** | **<0.001** | | |
| **Cohort mean FEC** | **-0.471** | **0.183** | **0.010** | | |
| **Birth weight** | **0.337** | **0.061** | **<0.001** | | |

| | | | |
|---|---|---|---|
| Sex (male):Natal litter size (twin) | 0.316 | 0.221 | 0.152 |
| Sex (male):Maternal loss (mother survived) | -0.039 | 0.255 | 0.877 |
| **Sex (male):Population size YOB** | **-0.312** | **0.125** | **0.013** |
| Sex (male):Gestational winter NAO | -0.047 | 0.093 | 0.609 |
| **Sex (male):First winter NAO** | **-0.296** | **0.085** | **<0.001** |
| **Sex (male):Cohort mean FEC** | **-0.213** | **0.101** | **0.035** |
| Sex (male):Birth weight | -0.038 | 0.087 | 0.660 |

B) Lifetime Breeding Success (N=1721)

| | | | | | |
|---|---|---|---|---|---|
| **Intercept** | **1.293** | **0.168** | **<0.001** | Maternal identity | 0.208 |
| **Sex (male)** | **-0.675** | **0.295** | **0.022** | Birth year | 0.127 |
| Natal litter size (twin) | 0.030 | 0.128 | 0.816 | | |
| Maternal loss (mother survived) | 0.289 | 0.158 | 0.068 | Dispersion | 0.932 |
| Population size YOB | 0.039 | 0.092 | 0.674 | | |
| First winter NAO | 0.016 | 0.079 | 0.840 | | |
| Cohort mean FEC | -0.009 | 0.083 | 0.914 | | |
| **Birth weight** | **0.155** | **0.045** | **0.001** | | |
| Sex (male):Natal litter size (twin) | 0.075 | 0.196 | 0.703 | | |
| Sex (male):Maternal loss (mother survived) | -0.065 | 0.301 | 0.828 | | |
| **Sex (male):Population size YOB** | **-0.183** | **0.084** | **0.029** | | |
| **Sex (male):First winter NAO** | **0.164** | **0.070** | **0.019** | | |
| Sex (male):Cohort mean FEC | -0.075 | 0.086 | 0.386 | | |
| **Sex (male):Birth weight** | **0.171** | **0.072** | **0.017** | | |

C) Longevity (N=1721)

| | | | | | |
|---|---|---|---|---|---|
| **Intercept** | **1.513** | **0.094** | **<0.001** | Maternal identity | 0.009 |
| **Sex (male)** | **-0.507** | **0.156** | **0.001** | Birth year | 0.067 |

| | | | | | |
|---|---|---|---|---|---|
| Natal litter size (twin) | -0.035 | 0.065 | 0.585 | | |
| **Maternal loss (mother survived)** | **0.180** | **0.082** | **0.028** | Dispersion | 4.69 |
| Population size YOB | 0.086 | 0.061 | 0.162 | | |
| First winter NAO | 0.041 | 0.053 | 0.439 | | |
| Cohort mean FEC | -0.051 | 0.054 | 0.350 | | |
| **Birth weight** | **0.079** | **0.023** | **0.001** | | |
| Sex (male):Natal litter size (twin) | -0.011 | 0.108 | 0.916 | | |
| Sex (male):Maternal loss (mother survived) | -0.241 | 0.160 | 0.133 | | |
| Sex (male):Population size YOB | 0.011 | 0.046 | 0.811 | | |
| **Sex (male):First winter NAO** | **0.121** | **0.039** | **0.002** | | |
| Sex (male):Cohort mean FEC | -0.013 | 0.047 | 0.774 | | |
| Sex (male):Birth weight | -0.049 | 0.039 | 0.210 | | |

D) Breeding probability (N=9572 of 2232 individuals)

| | | | | | |
|---|---|---|---|---|---|
| **Intercept** | **1.544** | **0.248** | **<0.001** | Identity | 1.529 |
| **Sex (male)** | **-2.605** | **0.405** | **<0.001** | Maternal identity | 0.337 |
| **Age (in years)** | **79.581** | **4.526** | **<0.001** | Measurement year | 0.318 |
| **Age (in years)$^2$** | **-107.651** | **4.394** | **<0.001** | Birth year | 0.144 |
| Natal litter size (twin) | -0.350 | 0.186 | 0.061 | | |
| **Maternal loss (mother survived)** | **0.551** | **0.225** | **0.015** | | |
| **Population size YOB** | **-0.288** | **0.112** | **0.010** | | |
| First winter NAO | -0.038 | 0.096 | 0.694 | | |
| Cohort mean FEC | -0.070 | 0.101 | 0.493 | | |
| **Birth weight** | **0.238** | **0.067** | **<0.001** | | |
| Sex (male):Age (in years) | -5.330 | 12.096 | 0.659 | | |
| Sex (male):Age (in years)$^2$ | 6.946 | 11.228 | 0.536 | | |
| Sex (male):Natal litter size (twin) | 0.409 | 0.300 | 0.173 | | |
| Sex (male):Maternal loss (mother survived) | -0.225 | 0.411 | 0.585 | | |

| | | | |
|---|---|---|---|
| **Sex (male):Population size YOB** | **-0.392** | **0.125** | **0.002** |
| Sex (male):First winter NAO | -0.144 | 0.104 | 0.165 |
| **Sex (male):Cohort mean FEC** | **0.323** | **0.128** | **0.012** |
| Sex (male):Birth weight | 0.057 | 0.106 | 0.591 |

Table S3. Fixed and random effect estimates from a negative binomial GLMM of longevity in Soay sheep that survived beyond the first year of life. Statistically significant fixed effects are highlighted in bold. The reference levels for the fixed factors were: sex (female), natal litter size (singleton) and maternal loss (mother died in first year of life). The fixed covariates were scaled to mean=0 and standard deviation=1. Dispersion parameter for negative binomial model 4.69.

| N=1721 | | | | | |
|---|---|---|---|---|---|
| Fixed effects | Estimate | Std. Error | p-value | Random effects | Variance |
| **Intercept** | **1.569** | **0.086** | **<0.001** | Maternal identity | 0.011 |
| **Sex (male)** | **-0.740** | **0.037** | **<0.001** | Birth year | 0.067 |
| Natal litter size (twin) | -0.041 | 0.053 | 0.441 | | |
| Maternal loss (mother survived) | 0.119 | 0.071 | 0.093 | | |
| Population size YOB | 0.088 | 0.060 | 0.144 | | |
| First winter NAO | 0.041 | 0.053 | 0.441 | | |
| Cohort mean FEC | -0.055 | 0.053 | 0.299 | | |
| **Birth weight** | **0.063** | **0.020** | **0.001** | | |
| **Sex (male):First winter NAO** | **0.120** | **0.036** | **0.001** | | |

Table S4. Fixed and random effect estimates from a binomial GLMM of annual breeding probability in Soay sheep that survived beyond the first year of life. Statistically significant fixed effects are highlighted in bold. The reference levels for the fixed factors were: sex (female), natal litter size (singleton) and maternal loss (mother died in first year of life). The fixed covariates were scaled to mean=0 and standard deviation=1.

| N=9572 of 2232 | | | | | |
|---|---|---|---|---|---|
| Fixed effects | Estimate | Std. Error | p-value | Random effects | Variance |
| **Intercept** | **1.576** | **0.221** | **<0.001** | Identity | 1.539 |
| **Sex (male)** | **-2.724** | **0.114** | **<0.001** | Maternal identity | 0.324 |
| **Age (in years)** | **77.910** | **4.215** | **<0.001** | Measurement year | 0.322 |
| **Age (in years)$^2$** | **-105.724** | **3.972** | **<0.001** | Birth year | 0.141 |
| Natal litter size (twin) | -0.201 | 0.152 | 0.187 | | |

| | | | |
|---|---|---|---|
| **Maternal loss (mother survived)** | **0.488** | **0.190** | **0.010** |
| **Population size YOB** | **-0.301** | **0.111** | **0.007** |
| First winter NAO | -0.086 | 0.090 | 0.341 |
| Cohort mean FEC | -0.056 | 0.101 | 0.575 |
| **Birth weight** | **0.260** | **0.054** | **<0.001** |
| **Sex (male):Population size YOB** | **-0.350** | **0.121** | **0.004** |
| **Sex (male):Cohort mean FEC** | **0.283** | **0.126** | **0.025** |

Table S5. Fixed and random effect estimates from a binomial GLMM of annual twinning probability for female Soay sheep that bred that year. Statistically significant fixed effects are highlighted in bold. The reference levels for the fixed factors were: natal litter size (singleton) and maternal loss (mother died in first year of life). The fixed covariates were scaled to mean=0 and standard deviation=1.

| N=5580 of 1092 | | | | | |
|---|---|---|---|---|---|
| Fixed effects | Estimate | Std. Error | p-value | Random effects | Variance |
| **Intercept** | **-2.871** | **0.314** | **<0.001** | Identity | 0.988 |
| **Age (in years)** | **79.973** | **5.277** | **<0.001** | Maternal identity | 0.982 |
| **Age (in years)$^2$** | **-39.741** | **4.561** | **<0.001** | Measurement year | 0.404 |
| **Natal litter size (twin)** | **0.702** | **0.246** | **0.004** | Birth year | <0.001 |
| Maternal loss (mother survived) | -0.332 | 0.307 | 0.280 | | |
| Population size YOB | -0.115 | 0.098 | 0.241 | | |
| First winter NAO | -0.073 | 0.078 | 0.348 | | |
| Cohort mean FEC | -0.021 | 0.091 | 0.821 | | |
| **Birth weight** | **0.301** | **0.081** | **<0.001** | | |

Table S6. Fixed and random effect estimates from a negative binomial GLMM of annual offspring number for male Soay sheep that bred that year. Statistically significant fixed effects are highlighted in bold. The reference levels for the fixed factors were: natal litter size (singleton) and maternal loss (mother died in first year of life). The fixed covariates were scaled to mean=0 and standard deviation=1. Dispersion parameter for negative binomial model 40.7.

| N=884 of 378 | | | | | |
|---|---|---|---|---|---|
| Fixed effects | Estimate | Std. Error | p-value | Random effects | Variance |
| **Intercept** | **0.449** | **0.161** | **0.005** | Identity | 0.146 |
| **Age (in years)** | **15.250** | **0.838** | **<0.001** | Maternal identity | 0.020 |

| | | | | | |
|---|---|---|---|---|---|
| **Age (in years)$^2$** | **-3.142** | **0.705** | **<0.001** | Measurement year | 0.028 |
| Natal litter size (twin) | 0.027 | 0.116 | 0.813 | Birth year | <0.001 |
| **Maternal loss (mother survived)** | **0.334** | **0.163** | **0.040** | | |
| **Population size YOB** | **-0.104** | **0.048** | **0.030** | | |
| First winter NAO | -0.045 | 0.037 | 0.227 | | |
| Cohort mean FEC | 0.037 | 0.045 | 0.412 | | |
| **Birth weight** | **0.125** | **0.038** | **0.001** | | |

Table S7. Fixed and random effect estimates from GLMMs of A) lifetime breeding success (negative binomial), B) longevity (negative binomial), C) breeding probability, D) female twinning probability, and E) male offspring number in Soay sheep, accounting for mean adult body weight (measured in August, corrected for age and sex). Statistically significant fixed effects are highlighted in bold. The reference levels for the fixed factors were: sex (female), natal litter size (singleton) and maternal loss (mother died in first year of life). The fixed covariates were scaled to mean=0 and standard deviation=1.

| **Fixed effects** | **Estimate** | **Std. Error** | **p-value** | **Random effects** | **Variance** |
|---|---|---|---|---|---|
| A) Lifetime Breeding Success (N=1263) | | | | | |
| **Intercept** | **1.468** | **0.132** | **<0.001** | Maternal identity | 0.129 |
| **Sex (male)** | **-0.470** | **0.073** | **<0.001** | Birth year | 0.034 |
| **Mean adult weight** | **0.292** | **0.037** | **<0.001** | | |
| Natal litter size (twin) | 0.083 | 0.097 | 0.394 | Dispersion | 1.37 |
| **Maternal loss (mother survived)** | **0.280** | **0.129** | **0.030** | | |
| **Population size YOB** | **0.121** | **0.061** | **0.048** | | |
| First winter NAO | 0.032 | 0.053 | 0.545 | | |
| Cohort mean FEC | -0.073 | 0.054 | 0.176 | | |
| Birth weight | 0.049 | 0.041 | 0.234 | | |
| Sex (male):Population size YOB | -0.087 | 0.069 | 0.209 | | |
| **Sex (male):First winter NAO** | **0.147** | **0.066** | **0.026** | | |
| **Sex (male): Birth weight** | **0.215** | **0.067** | **0.001** | | |
| B) Longevity (N=1263) | | | | | |
| **Intercept** | **1.707** | **0.075** | **<0.001** | Maternal identity | 0.007 |
| **Sex (male)** | **-0.713** | **0.038** | **<0.001** | Birth year | 0.029 |

| | | | | | |
|---|---|---|---|---|---|
| **Mean adult weight** | **0.128** | **0.019** | **<0.001** | | |
| Natal litter size (twin) | -0.043 | 0.051 | 0.400 | Dispersion | 9.34 |
| Maternal loss (mother survived) | 0.133 | 0.068 | 0.051 | | |
| **Population size YOB** | **0.139** | **0.043** | **0.001** | | |
| First winter NAO | 0.041 | 0.038 | 0.271 | | |
| Cohort mean FEC | -0.073 | 0.039 | 0.057 | | |
| Birth weight | 0.019 | 0.019 | 0.306 | | |
| **Sex (male):First winter NAO** | **0.133** | **0.037** | **<0.001** | | |

C) Breeding probability (N=8034 of 1491 individuals)

| | | | | | |
|---|---|---|---|---|---|
| **Intercept** | **2.120** | **0.240** | **<0.001** | Identity | 1.407 |
| **Sex (male)** | **-2.532** | **0.127** | **<0.001** | Maternal identity | 0.291 |
| Mean adult weight | 0.066 | 0.053 | 0.211 | Measurement year | 0.307 |
| **Age (in years)** | **56.694** | **3.903** | **<0.001** | Birth year | 0.046 |
| **Age (in years)$^2$** | **-98.157** | **3.896** | **<0.001** | | |
| Natal litter size (twin) | -0.100 | 0.168 | 0.552 | | |
| Maternal loss (mother survived) | 0.227 | 0.220 | 0.302 | | |
| **Population size YOB** | **-0.259** | **0.098** | **0.008** | | |
| First winter NAO | -0.055 | 0.073 | 0.455 | | |
| Cohort mean FEC | -0.084 | 0.088 | 0.340 | | |
| **Birth weight** | **0.185** | **0.062** | **0.003** | | |
| Sex (male):Population size YOB | -0.270 | 0.140 | 0.054 | | |
| Sex (male):Cohort mean FEC | 0.220 | 0.147 | 0.134 | | |

D) Female Twinning probability (N=5232 of 934 individuals)

| | | | | | |
|---|---|---|---|---|---|
| **Intercept** | **-2.807** | **0.315** | **<0.001** | Identity | 0.809 |
| **Mean adult weight** | **0.606** | **0.084** | **<0.001** | Maternal identity | 0.953 |
| **Age (in years)** | **74.352** | **5.175** | **<0.001** | Measurement year | 0.164 |
| **Age (in years)$^2$** | **-39.650** | **4.529** | **<0.001** | Birth year | <0.001 |
| **Natal litter size (twin)** | **0.775** | **0.247** | **0.002** | | |

| | | | |
|---|---|---|---|
| Maternal loss (mother survived) | -0.382 | 0.309 | 0.216 |
| Population size YOB | -0.065 | 0.099 | 0.510 |
| First winter NAO | -0.050 | 0.079 | 0.525 |
| Cohort mean FEC | -0.037 | 0.093 | 0.693 |
| Birth weight | 0.153 | 0.085 | 0.070 |

E) Male offspring number (N=727 of 278 individuals)

| | | | | | |
|---|---|---|---|---|---|
| **Intercept** | **0.500** | **0.186** | **0.007** | Identity | 0.162 |
| **Mean adult weight** | **0.194** | **0.043** | **<0.001** | Maternal identity | <0.001 |
| **Age (in years)** | **13.806** | **0.821** | **<0.001** | Measurement year | 0.035 |
| **Age (in years)$^2$** | **-3.130** | **0.691** | **<0.001** | Birth year | <0.001 |
| Natal litter size (twin) | 0.125 | 0.128 | 0.327 | | |
| Maternal loss (mother survived) | 0.347 | 0.188 | 0.065 | Dispersion | 38.8 |
| Population size YOB | -0.055 | 0.054 | 0.307 | | |
| First winter NAO | -0.059 | 0.041 | 0.150 | | |
| Cohort mean FEC | 0.042 | 0.051 | 0.407 | | |
| **Birth weight** | **0.099** | **0.044** | **0.025** | | |

Figure S1. Associations between measures of early-life adversity in Soay sheep. Pearson's moment correlation coefficients shown in Table S1. Points show the raw data and/or cohort means.

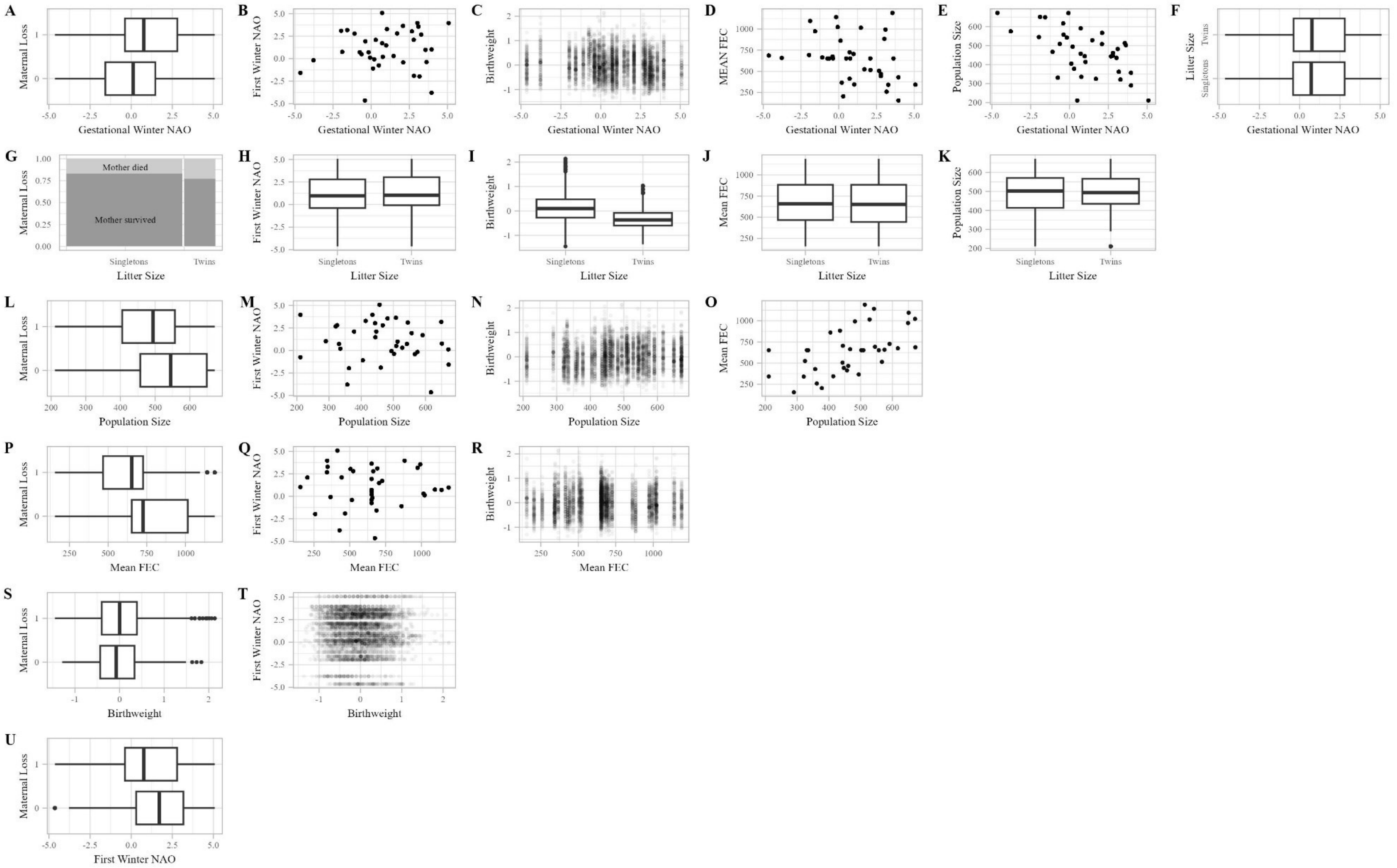
